# So ManyFolds, So Little Time: Efficient Protein Structure Prediction With pLMs and MSAs

**DOI:** 10.1101/2022.10.15.511553

**Authors:** Thomas D. Barrett, Amelia Villegas-Morcillo, Louis Robinson, Benoit Gaujac, David Adméte, Elia Saquand, Karim Beguir, Arthur Flajolet

## Abstract

In recent years, machine learning approaches for *de novo* protein structure prediction have made significant progress, culminating in AlphaFold which approaches experimental accuracies in certain settings and heralds the possibility of rapid *in silico* protein modelling and design. However, such applications can be challenging in practice due to the significant compute required for training and inference of such models, and their strong reliance on the evolutionary information contained in multiple sequence alignments (MSAs), which may not be available for certain targets of interest. Here, we first present a streamlined AlphaFold architecture and training pipeline that still provides good performance with significantly reduced computational burden. Aligned with recent approaches such as OmegaFold and ESMFold, our model is initially trained to predict structure from sequences alone by leveraging embeddings from the pretrained ESM-2 protein language model (pLM). We then compare this approach to an equivalent model trained on MSA-profile information only, and find that the latter still provides a performance boost – suggesting that even state-of-the-art pLMs cannot yet easily replace the evolutionary information of homologous sequences. Finally, we train a model that can make predictions from either the combination, or only one, of pLM and MSA inputs. Ultimately, we obtain accuracies in any of these three input modes similar to models trained uniquely in that setting, whilst also demonstrating that these modalities are complimentary, each regularly outperforming the other.

## 1 Introduction

Proteins are the building blocks of all cellular life. Understanding their 3D structure is essential to understanding their function and, in principle, these structures are predictable from only the amino acid sequence [1, 2]. However, in practice this is highly challenging due to the complex many-body atomic interactions. In recent years, deep learning based approaches have made significant strides in protein folding – with recent models such as AlphaFold [3] and RoseTTAFold [4] approaching experimental accuracies in certain settings and heralding a new era of structurally-rich bioinformatics.

However, this remarkable performance is delivered by a considerable computational workload during training and inference, that makes both deployment and further model development practically challenging. Moreover, despite these costs, current state-of-the-art approaches do not predict structure from sequence alone. For example, AlphaFold is heavily reliant on large multiple sequence alignments (MSAs) and, to a lesser degree, templates of similar sequences with known structure. Indeed, generating sufficient MSAs is an extensive process requiring searching though large structural databases, and, even then, these can be of low-quality for rare proteins that lack known homologs [5].

Within the last year, an alternative to MSA-based models has emerged – leveraging information contained in pre-trained protein language models (pLMs) [6, 7]. These pLMs are trained on vast sequence databases in a self-supervised manner (with the objective of predicting labels for masked amino acids in a protein sequence). The internal representations (embeddings) learned have proven successful for many downstream tasks, including predicting structural attributes of the protein [6–11]. An appealing approach is therefore to replace explicit representations of external sequence/structural databases (i.e. MSAs/templates) with pLM embeddings, which has been the foundation of recent models that predict 3D structures from protein sequence alone [5, 12, 13].

In this work, we begin by presenting a streamlined AlphaFold-like model (MonoFold) that offers faster inference and can be trained to a reasonably strong level of performance within a more limited compute budget (∼1 day on a v2-128 Google TPU). Using this, we train different models that take in either pLM embeddings (from the state-of-the-art ESM-2 pLM [12]) or a compressed statistical representation of the MSA. We find that even our smaller model with a reduced training budget can approach or outperform the accuracy of existing pLM-based approaches. Moreover, using the target MSA-profile provides a consistent boost in performance over our pLM-only model, which suggests that whilst protein embeddings do contain relevant structural information, they can not immediately replace explicit search for homologous sequences without cost.

Next, we consider the potential synergies between pLM and MSA information by training a model that takes both as input (PolyFold). Initially, this model achieves similar performance to the MSA-only model previously trained, from which it could be interpreted that the pLM embeddings are not useful. However, after an additional targeted fine-tuning, our PolyFold model is able to predict structures using only the pLM or MSA inputs as accurately as the equivalent MonoFold variants, without loss of performance in the pLM+MSA setting. In addition to the increased flexibility this provides, we also find that different targets are better predicted using different inference modalities, with none of the three options (pLM-only, MSA-only, pLM+MSA) either dominating or being uniformly inferior. Ultimately, this underlines the utility of our unified model and opens the door to further pushing the limits of performance with multiple complimentary sources of input information.

## 2 MonoFold and PolyFold models

The full AlphaFold model takes as input the raw sequence, MSAs – both clustered and additional “extra” raw MSA sequences – and, optionally, templates of homologous proteins with resolved structures. From these, 1D (per residue position with one set per MSA cluster) and 2D (between pairwise residues) embeddings, are extracted and iteratively updated by a stack of Evoformer modules. Finally, the Structure Module takes the first row of the 1D features – dubbed the “single representation” – and the 2D “pair representation” to predict a final 3D geometry for the full protein.

Our models, henceforth called MonoFold and PolyFold for simplicity, remove the template and “extra” MSA components of the original AlphaFold entirely. The input information is modified to include one (MonoFold), or both (PolyFold), of pLM embeddings and an MSA profile. As well as reducing the dimension of the 1D processing track to match these modified inputs, the Evoformer is also made less computationally burdensome by removing or streamlining certain expensive operations. This section highlights key modifications, with further details of our input pre-processing pipeline and architectural changes in Appendix A. Architectural and data pipeline details that are unchanged from AlphaFold are provided in [3].

### 2.1 Input features

#### pLM embeddings

We used ESM-2 with 650M parameters [12] as our pLM. Similarly to ESMFold [12], we projected the output per-residue embeddings to obtain our 1D features. The 2D pair representation is initialised with the pairwise relative positional encoding of AlphaFold [3], plus the projection of the attention maps from each hidden layer of the ESM-2.

#### MSA statistics

To maintain a lightweight architecture with the same input dimensions as in the pLM case, we choose not to stack information from multiple MSA clusters. Instead, we follow an approach closely inspired by the “no raw MSA” ablation of AlphaFold (detailed in SM 1.13 of [3] and shown to provide only a small performance degradation). Concretely, our 1D input is the equivalent to setting the number of MSA clusters to 1, with the cluster centre being the target sequence. In addition to the pairwise relative positional encoding, the pair representation is initialised by projecting 1024 raw “extra MSA” features, followed by an `OuterProductMean` module (Alg. 10 in SM of [3]).

#### Combined pLM and MSA input

We also train a model, PolyFold, that combines both the pLM embeddings and MSA statistics detailed above into a single input. As the only the first row of the 1D features are passed from the Evoformer to the Structure Module, we found that the model was biased by which input features we set as the first row. Therefore, we instead initialise the 1D features by stacking a projection of the one-hot encoded target sequence (as the first row) with the 1D pLM and MSA inputs. The 2D pair representation is initialised by combining both the pLM and MSA 2D initialisations with the pairwise relative positional encoding.

### 2.2 Evoformer modifications

Our architecture is summarised in Figure 1, which details the operations contained in the modified Evoformer. The Evoformer takes in 1D and 2D features, and returns updated features with unchanged dimensions following a series of communication, attention and transition modules. The details of these operations are unchanged to those presented in [3] with a few key exceptions.

**Figure 1:**
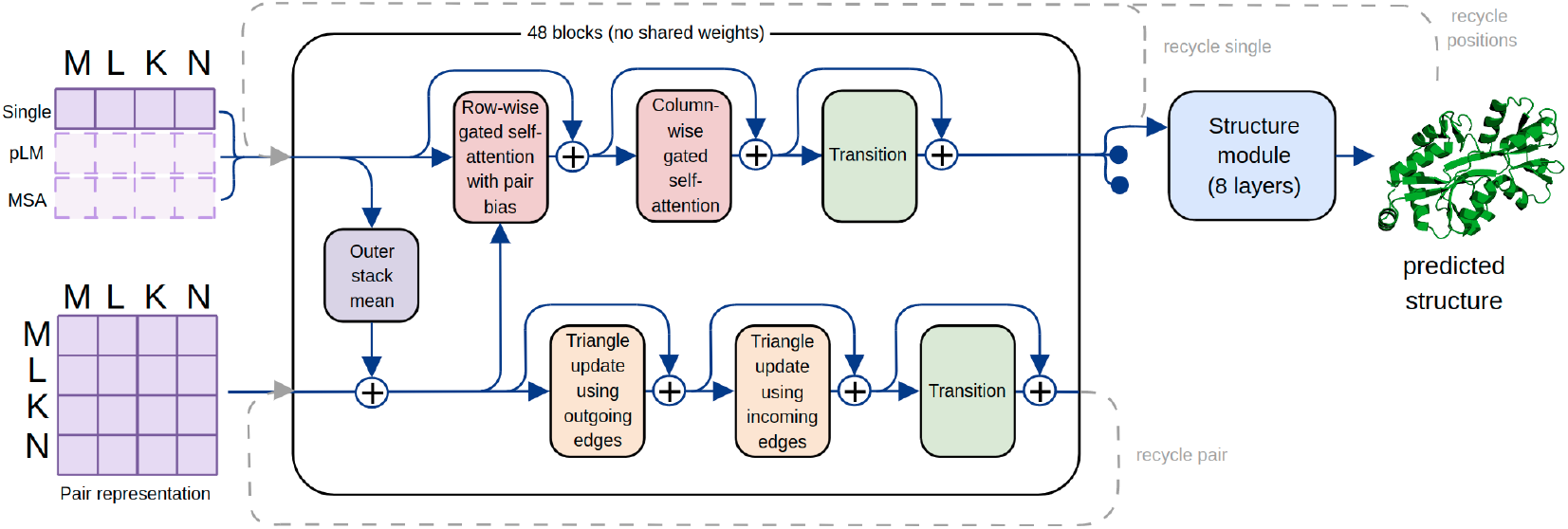
The summarised architecture of our PolyFold model. For MonoFold, only one of the input rows is used and the column-wise attention in the upper track is removed.

Firstly, the communication from the 1D track to the 2D track is moved to the beginning of the block – as was done for AlphaFold Multimer [14] to reduce the memory requirements for training – and is performed using a custom `OuterStackMean` operation, replacing an outer product operation with a cheaper concatenation operation (see Appendix Alg. 1). Secondly, we remove the two “triangular self-attention” modules used to process the pair representation in the full AlphaFold model, retaining only the “triangular update” modules, as the former are especially expensive during training and inference. We also note that column-wise attention module is only meaningful (and, therefore, only included) when processing the joint pLM and MSA features, to attend across all three input rows.

## 3 Experimentss

### Training details

We trained our models from scratch using three AlphaFold losses – the structure module loss (weight 1), distogram loss (weight 0.3), and pLDDT loss (weight 0.01) – and a batch sizes of 128. When a pLM is used, samples are first uniformly cropped (or padded) to a contiguous section of 1024 residues, before being processed through ESM-2 – which is frozen throughout training – and then subsequently cropped to a reduced length of 256 residues before being passed to the folding model. Note that we did not perform any fine-tuning on larger crops or deploy self-distillation on extended datasets of predicted structures. We use an Adam optimizer [15] with default parameters and learning rate of (10^−3^, with a linear warm-up of 1000 steps. Each model was initially trained for 20 k steps on Google TPUs v2-128 (one sample per core), which is approximately 25-29 hours of wall-clock time depending on the model. To run our experiments, we extended our Jax-based library for training AlphaFold and related models [16], which is publicly available at https://github.com/instadeepai/manyfold.

### Validation and metrics

The models were validated on exponentially weighted averaged (EWA) parameters using a decay of 0.999. During inference, we used 3 recycling iterations. As performance metrics, we considered the standard lDDT score [17] (0-1) and TM-score [18] (0-100), widely used in previous works and CASP challenges. We run the validation inference on an NVIDIA A100 GPU.

### Datasets

The training set consists of entries in the Protein Data Bank (PDB) [19] with a release date before 2020-05-14, a resolution *<* 9Å, and no single amino acid accounting for more than 80% of the sequence. This adds up to approximately 490 k structures. During training the same stochastic filters as in [3] were used to sample each batch.

To validate our models, we used targets from the CAMEO [20] and CASP14 [21] competitions. For the former, we collected 143 CAMEO targets released from March to May 2022, with less than 700 residues. This includes samples of the three levels of difficulty (easy, medium and hard). For the latter, we extracted the domain-level targets from the Free-Modeling (FM) and Template-Based Modeling hard (TBM-hard) categories of CASP14, only considering contiguous domains that are part of protein chains added to the PDB. This results in 34 target domains with a maximum sequence length of 405 residues. The full list of validation targets can be found in Appendix B.

### Baselines

Our primary baseline is AlphaFold (`model_1_ptm`), which utilises both MSAs and templates, as provided by the offical release code. We also run AlphaFold “sequence-only”, i.e. without MSAs or templates (`model_5_ptm` zeroing and masking MSA features). We compare to recent pLM-based models trained for protein structure prediction from sequence alone – specifically OmegaFold [5] and HelixFold-Single [13], in both cases using the official publicly available code and checkpoints. The recent ESMFold [12] model (which is a minimally modified AlphaFold Evoformer and Structure Module operating on ESM-2 embeddings) does not yet have a public release, and so we cannot report a direct comparison given the different validation sets between their paper and ours. However, as detailed below, we can still infer the approximate relative performance of our models from the reported results and do so to provide context, rather than as an absolute comparison.

### 3.1 Inference and training timings

To assess the efficiency of our models, we measure the inference and training timings of our PolyFold model compared to AlphaFold `model_1_ptm`. For inference, we run forward passes on a NVIDIA A100 GPU of individual samples with lengths varying from 100 to 700 residues, and average the resulting times at each length over 5 samples. Figure 2 shows the inference timings, where we see that our PolyFold model scales better with the sequence length, reaching *×*6 time reduction with respect to AlphaFold for the longer lengths. For a full comparison, here we also include the inference times we obtain for OmegaFold, using its official implementation [5]. We see that the OmegaFold model is slightly faster than AlphaFold for shorter sequences, but scales worse with sequence length. To measure the training step times, we use batches of 128 samples with a crop size of 256. We run AlphaFold on TPUs v3-8 and PolyFold on TPUs v2-128, obtaining approximately 13.9 seconds for AlphaFold and 4.7 seconds for PolyFold. This reduction in both training time and memory load makes our models more efficient to train and validate, allowing more experiments and analyses to be performed in less time, while requiring fewer computing resources.

**Figure 2:**
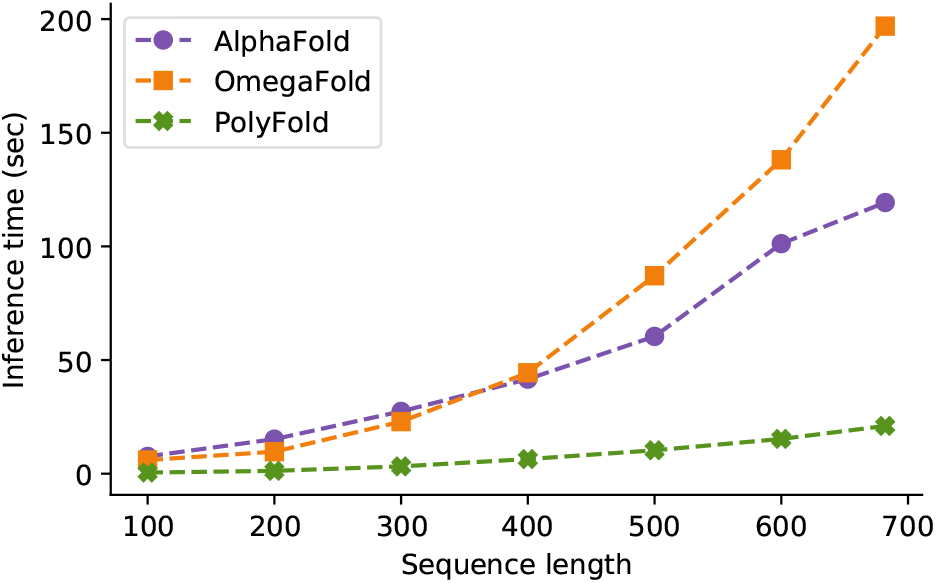
Inference timings measured for different input sequence lengths on the CAMEO dataset.

### 3.2 Single-input models: pLMs versus MSAs

The performance of models trained (from scratch for 20 k steps) on only ESM-2 embeddings or MSA-profile statistics are shown in Figure 3. We first note that even with this limited training budget, these compressed models still provide quite strong performance, with TM-scores of 79.5 and 82.6 on the CAMEO dataset (thus achieving 89 % and 93 % of the performance of AlphaFold).

**Figure 3:**
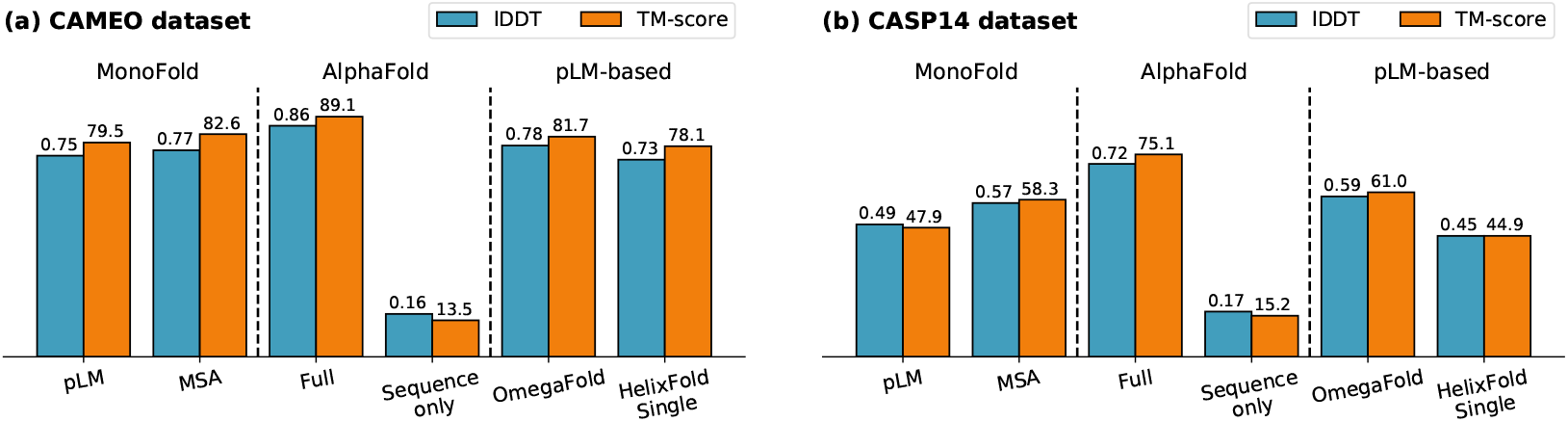
Performance of MonoFold, with pLM or MSA inputs, on (a) CAMEO and (b) CASP14 datasets. Baselines include AlphaFold and recent pLM-based folding models.

Indeed, our MonoFold-pLM model outperforms HelixFold-Single and is already approaching the performance of OmegaFold. As our training budget is not sufficient to reach convergence, this gap would likely close further, though it is unclear whether it would be expected to match or exceed OmegaFold (which has a different architecture and pLM). Whilst we can not directly compare to ESMFold, the original paper [12] reports a TM-score of 82.8 on a CAMEO dataset of similar difficulty (as judged by AlphaFold’s full strength performance of 88.3 and 89.1 on their and our dataset, respectively). This suggests that MonoFold-pLM is approaching, but below, the accuracy of ESMFold, which is unsuprising as ESMFold has a larger Evoformer and training budget.

Notably, MonoFold-MSA is even better, already matching the strongest pLM-only models whilst again not being trained to convergence. This suggests that MSAs still appear to be more informative even with our considered ESM-2 model, and so pLM’s can not yet be used as a complete replacement for the evolutionary information of homolgous sequences.

The CASP14 dataset is more challenging for all models and, whilst this further emphasises the difference between models, the relative comparisons and analysis remain largely unchanged.

### 3.3 Combining pLM embeddings and MSA-profile

#### Combined pLM and MSA input

We next train PolyFold, which takes both ESM-2 embeddings and MSA-profile statistics as input. The performance after 20 k training steps are shown in the left columns of Figure 4 where we see that this combined model has essentially the same performance as MonoFold-MSA. This suggests that the information in the pLM and MSAs are not jointly more informative than MSAs alone, albeit training to convergence may reveal a difference in top-end performance. However, when either of the inputs is removed (by zeroing and masking the features), the performance significantly drops, so pLM embeddings are not simply being ignored.

**Figure 4:**
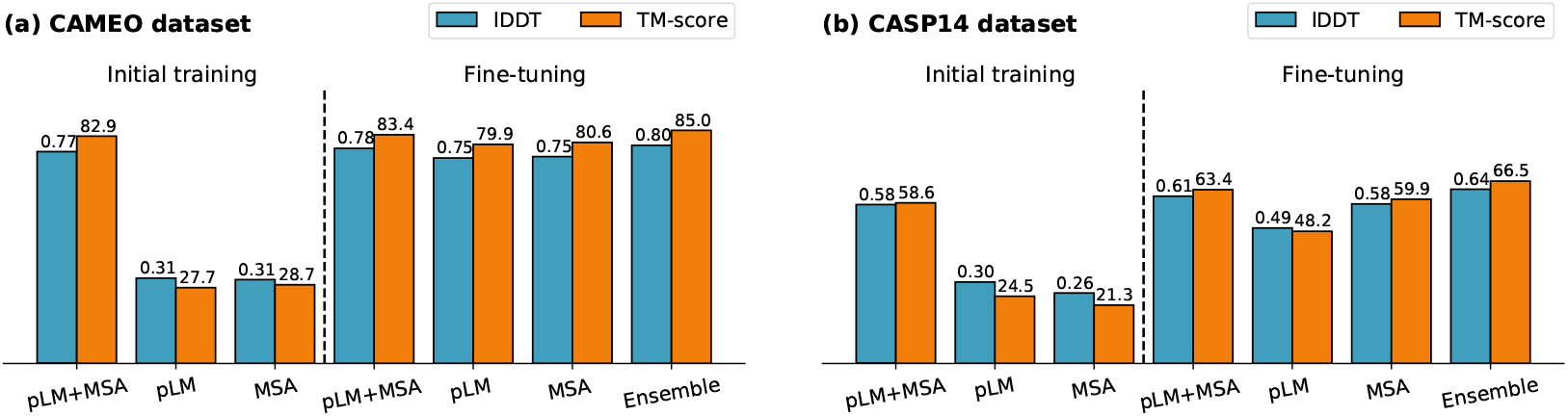
Performance of PolyFold, with pLM+MSA, pLM-only and MSA-only inference. After fine-tuning for strong performance across all inference modes, the “Ensemble” performance – where the best score from each mode is used – significantly outperforms full pLM+MSA inference.

#### Fine-tuning for optional input

To further investigate the different operating modes of PolyFold, we fine-tuned the model for a further 20 k training steps while masking either the MSA-profile, the pLM embeddings, or neither (with equal probability). Validation results are summarised in the right columns of Figure 4. We see that the pLM+MSA accuracy is only slightly improved, however, inference with MSA- or pLM-only inputs is significantly more performant – approaching, or reaching, the performance of MonoFold models trained specifically in just these settings.

Besides the practical utility of being able to train and run a single model in multiple modalities depending on the available information, it is also interesting to consider how the performance of varies between these (recalling, for example, that one motivation for pLM-based models is that MSAs can be difficult to obtain for certain sequences). Strikingly, we find the single-masked-input inference modes regularly outperform the full-information inference, with MSA-only (pLM-only) inference providing better TM-scores than pLM+MSA inference on 34.3 % (31.5 %) of the CAMEO validation set. Moreover, the MSA- and pLM-only modes are themselves better on different targets – if we always the score the best performing inference-mode on the CAMEO dataset, the overall TM-score rises to 85.0 with contributions split 25 %:32 %:43 % across pLM:MSA:pLM+MSA modes.

## 4 Discussion

The ability to train performant protein folding models with reduced computational burden is an important step towards the efficient development of next-generation models. The specific directions explored in this work provide already insights and results in this direction. The clear utility of, even a compressed statistical representation of MSAs, in comparison to (and in combination with) pLM embeddings further underlines that even state-of-the-art foundational biology models such as ESM-2 cannot readily replace a more explicit representation of homologous sequences.

However, with pLM-based and MSA-based models each having unique advantages and disadvantages, we believe that architectures able to operate in multiple modes can provide a powerful tool adaptable to specific settings (e.g. the low-MSA settings of orphan or fast-evolving proteins). Moreover, generating multiple structure predictions for a single target, whether informed by different biases in the inputs such as in our case or otherwise, can further provide ensembling benefits and flexibility to practitioners using deep learning models. In this light, additional exploration of whether different inference modalities are indeed preferable in identifiable regimes (e.g. antibodies) and to what degree the ensembling performance improvement is related to a variety of input-information settings, as opposed to natural variation across models, are interesting avenues for further analysis that we presently defer to ongoing and future work.

## Acknowledgments

This research was supported with Cloud TPUs from Google’s TPU Research Cloud (TRC).

## A Model architecture

We detail here the modifications done in our models with respect to AlphaFold. For the remaining unchanged details see [3].

### A.1 Input pre-processing

#### A.1.1 1D input features

##### pLM features

We use the 650M parameters ESM-2 [12] with 33 transformer layers, embedding dimension of 1280 and 40 attention heads trained on UniRef50 [22]. As in ESMFold, we learn a weighted sum of the pLM embeddings for each layer as part of the folding model (Algorithm 2, SM 1.3 of [12]) producing a 1024-dimensional embedding feature.

##### MSA features

Similarly to the “no raw MSA” ablation of AlphaFold (SM 1.13 of [3]), we by-pass the MSA clustering pre-processing step, which is equivalent to setting the number of clusters to 1 and then concatenating the target sequence, the MSA-profile, and additional MSA features into a 49-dimensional representation (see Table 1 in [3]). The MSA-profile is defined as in SM 1.13 of [3] and corresponds to the empirical distribution of amino acids for each position in the sequence.

**Table 1:** The table is calculated with respect to the AlphaFold repository v2.2.4 in the low-memory setting, *si* are the sub-batching factors which are set to *s*_1_ = 4, *s*_2_ = 128. Note that r(*n*) is the complexity of evaluating a dummy variable axis in einsum, which can be treated as a reduce operation which can theoretically be computed in log(*n*) time.

| Module | Memory $\mathcal{O}(\cdot)$ | Time $\mathcal{O}(\cdot)$ |
| --- | --- | --- |
| MSARowAttnWithPairBias | $s_1 \cdot N_{\text{res}}^2 + N_{\text{msa}} \cdot N_{\text{res}}$ | $\frac{1}{s_1} N_{\text{msa}} \cdot r(N_{\text{res}})$ |
| MSAColumnAttention | $N_{\text{msa}}^2 \cdot s_1 + N_{\text{msa}} \cdot N_{\text{res}}$ | $\frac{1}{s_1} r(N_{\text{msa}}) \cdot N_{\text{res}}$ |
| OuterProdMean | $N_{\text{msa}} \cdot N_{\text{res}} + N_{\text{res}}^2$ | $\frac{1}{s_2} r(N_{\text{msa}}) \cdot N_{\text{res}}$ |
| TriangleUpdate | $N_{\text{res}}^2$ | $r(N_{\text{res}})$ |
| TriangleSelfAttn | $s_1 \cdot N_{\text{res}}^2$ | $\frac{1}{s_1} \cdot r(N_{\text{res}}) \cdot N_{\text{res}}$ |
| MSAColumnGlobalAttn | $N_{\text{extra\_msa}} \cdot N_{\text{res}}$ | $\frac{1}{s_1} r(N_{\text{extra\_msa}}) \cdot N_{\text{res}}$ |
| ExtraMsa-MSARowAttnWPB | $s_1 \cdot N_{\text{res}}^2 + N_{\text{extra\_msa}} \cdot N_{\text{res}}$ | $\frac{1}{s_1} N_{\text{extra\_msa}} \cdot r(N_{\text{res}})$ |
| ExtraMsa-OuterProdMean | $N_{\text{extra\_msa}} \cdot N_{\text{res}} + N_{\text{res}}^2$ | $\frac{1}{s_2} r(N_{\text{extra\_msa}}) \cdot N_{\text{res}}$ |

##### 1D projection

In both cases, we project the 1D input representations using a 2-layer MLP with hidden dimension of 2056 and output dimension of 256.

#### A.1.2 2D input features

##### pLM features

As in the experiments without folding block in ESMFold [12], we extract and linearly project the pLM attention maps to initialise the pair representation. The projection produces a 128-dimensional pair feature.

##### MSA features

We use the extra MSA to initialise the pair representation in this case. Specifically, we first project 1024 extra MSA with a linear layer of size 128, then use an `OuterProductMean` module as described in Algorithm 10, SM 1.10 of [3]. We finally apply a dropout operation resulting in the pair representation of size 128.

##### pLM + MSA features

When using both pLM and MSA, we also combine the previous 2D representation. More specifically, we add the pLM pair representation and MSA pair representation using residual connections and a dropout layer, resulting in the 128-dimensional pair representation that is then fed to the Evoformer stack.

##### Relative position initialisation

On top of the 2D input representation, we include the pairwise relative positional encodings described in Algorithm 4 of [3], linearly projected to dimension 128.

### A.2 Evoformer

Inference with the full AlphaFold model and CASP14 hyperparameters is particularly expensive in memory and time. Not only do we want a faster inference time, but to perform a larger set of experiments we make changes also aimed at decreasing the training step time. Motivated by the expensive forward pass of AlphaFold [3], we chose to strip out the expensive modules which did not seem to cause a significant drop in performance when ablated. One particularly expensive module is the `TriangleAttention` in the 2D track, with slow execution time and a fairly large shard size of activations in memory (see Table 1). In the MonoFold models, we either use pLM embeddings or MSA statistics by setting the number of clusters to 1. This results in a single row for the 1D track, therefore reducing the size of the original MSA track from AlphaFold. Having only one row, we also remove the redundant column-wise attention operation for these models. The PolyFold model has three rows for the 1D track, so we keep the column-wise attention in this case. To reduce memory usage (to fit with 8 GB v2 TPU codes), in the PolyFold model the number of heads in the column- and row-attention modules of the 1D track is reduced from 8 to 4. Finally, for all models, we replace the `OuterProductMean` with an `Oute`rStackMean (see Algorithm 1). As we only have either one or three rows in the 1D track, we do not need to apply sub-batching and can keep the time complexity 𝒪 (*N*_msa_) = 𝒪 (1) also considering *N*_*res*_. For the rest of hyperparameters in the Evoformer modules refer to [3].

#### Algorithm 1 OuterStackMean

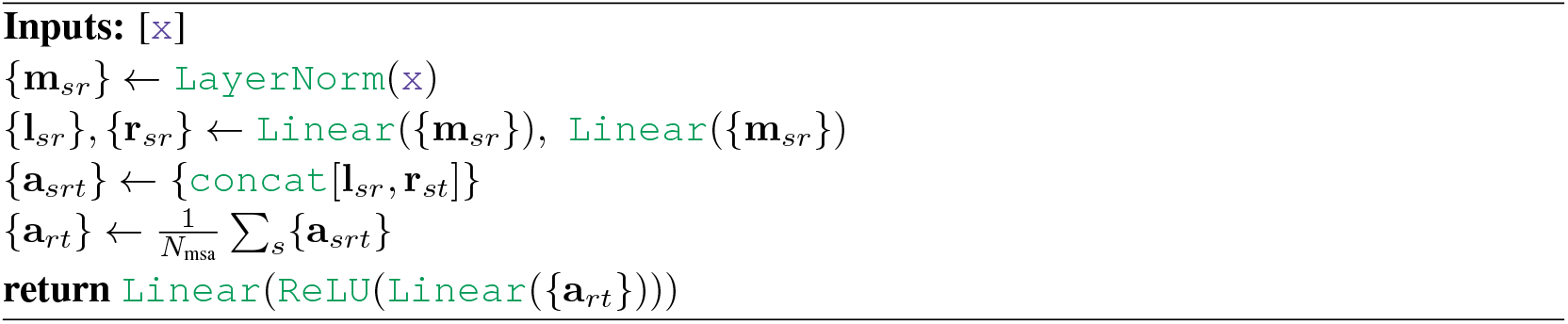

#### Algorithm 2 Inference

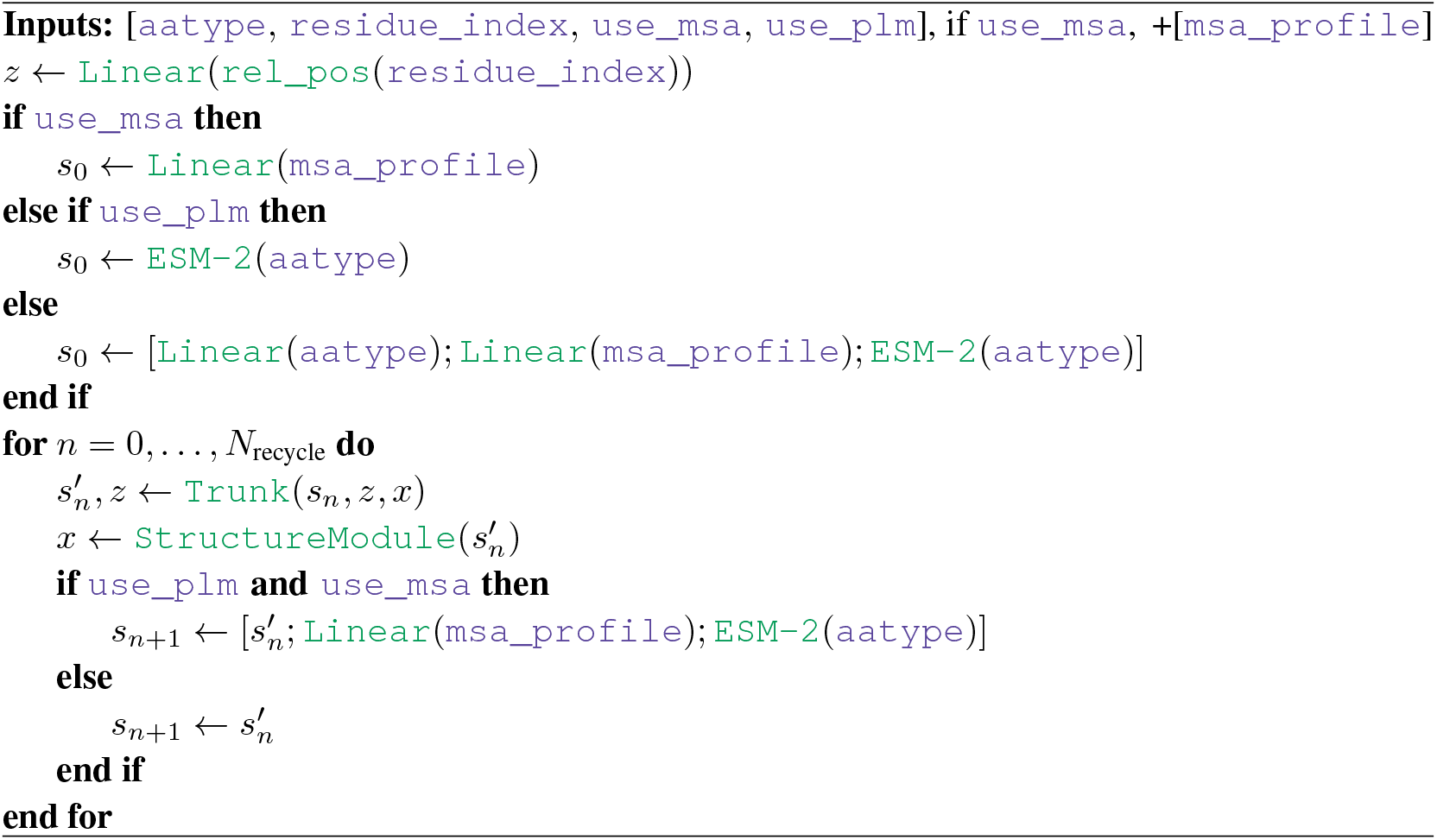

## B Validation targets

### B.1 CAMEO

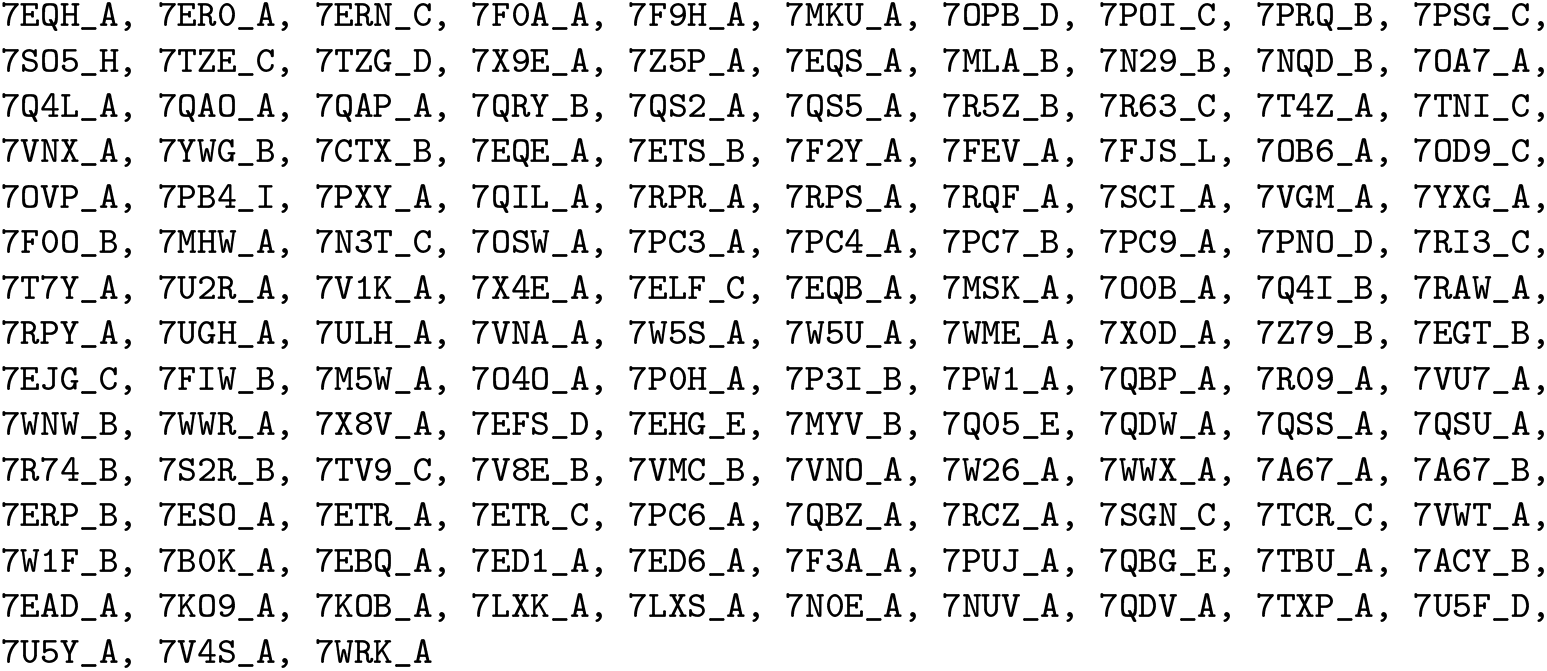

### B.2 CASP14

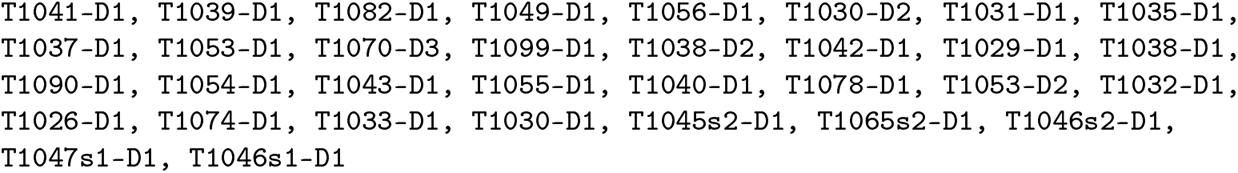

## References

[1] Christian B. Anfinsen. Principles that govern the folding of protein chains. Science, 181(4096): 223–230, 1973.

[2] Ken A. Dill, S. Banu Ozkan, M. Scott Shell, and Thomas R. Weikl. The protein folding problem. Annual Review of Biophysics, 37:289, 2008.

[3] John Jumper, Richard Evans, Alexander Pritzel, Tim Green, Michael Figurnov, Olaf Ronneberger, Kathryn Tunyasuvunakool, Russ Bates, Augustin Žídek, Anna Potapenko, et al. Highly accurate protein structure prediction with AlphaFold. Nature, 596(7873):583–589, 2021.

[4] Minkyung Baek, Frank DiMaio, Ivan Anishchenko, Justas Dauparas, Sergey Ovchinnikov, Gyu Rie Lee, Jue Wang, Qian Cong, Lisa N. Kinch, R. Dustin Schaeffer, et al. Accurate prediction of protein structures and interactions using a three-track neural network. Science, 373(6557):871–876, 2021.

[5] Ruidong Wu, Fan Ding, Rui Wang, Rui Shen, Xiwen Zhang, Shitong Luo, Chenpeng Su, Zuofan Wu, Qi Xie, Bonnie Berger, Jianzhu Ma, and Jian Peng. High-resolution de novo structure prediction from primary sequence. bioRxiv, 2022. doi: 10.1101/2022.07.21.500999.

[6] Alexander Rives, Joshua Meier, Tom Sercu, Siddharth Goyal, Zeming Lin, Jason Liu, Demi Guo, Myle Ott, C. Lawrence Zitnick, Jerry Ma, et al. Biological structure and function emerge from scaling unsupervised learning to 250 million protein sequences. Proceedings of the National Academy of Sciences, 118(15):e2016239118, 2021.

[7] Ahmed Elnaggar, Michael Heinzinger, Christian Dallago, Ghalia Rehawi, Wang Yu, Llion Jones, Tom Gibbs, Tamas Feher, Christoph Angerer, Martin Steinegger, Debsindhu Bhowmik, and Burkhard Rost. ProtTrans: Towards cracking the language of life’s code through self-supervised deep learning and high performance computing. IEEE Transactions on Pattern Analysis and Machine Intelligence, pages 1–16, 2021.

[8] Roshan Rao, Nicholas Bhattacharya, Neil Thomas, Yan Duan, Xi Chen, John Canny, Pieter Abbeel, and Yun S. Song. Evaluating protein transfer learning with TAPE. In Advances in Neural Information Processing Systems, 2019.

[9] Jesse Vig, Ali Madani, Lav R. Varshney, Caiming Xiong, Richard Socher, and Nazneen Rajani. BERTology meets biology: Interpreting attention in protein language models. In International Conference on Learning Representations, 2021.

[10] Brian Hie, Ellen D. Zhong, Bonnie Berger, and Bryan Bryson. Learning the language of viral evolution and escape. Science, 2021.

[11] Ratul Chowdhury, Nazim Bouatta, Surojit Biswas, Charlotte Rochereau, George M. Church, Peter K. Sorger, and Mohammed AlQuraishi. Single-sequence protein structure prediction using language models from deep learning. bioRxiv, 2021.

[12] Zeming Lin, Halil Akin, Roshan Rao, Brian Hie, Zhongkai Zhu, Wenting Lu, Allan dos Santos Costa, Maryam Fazel-Zarandi, Tom Sercu, Sal Candido, and Alexander Rives. Language models of protein sequences at the scale of evolution enable accurate structure prediction. bioRxiv, 2022. doi: 10.1101/2022.07.20.500902.

[13] Xiaomin Fang, Fan Wang, Lihang Liu, Jingzhou He, Dayong Lin, Yingfei Xiang, Xiaonan Zhang, Hua Wu, Hui Li, and Le Song. HelixFold-Single: MSA-free protein structure prediction by using protein language model as an alternative. arXiv, 2022. doi: 10.48550/ARXIV.2207.13921.

[14] Richard Evans, Michael O’Neill, Alexander Pritzel, Natasha Antropova, Andrew Senior, Tim Green, Augustin Žídek, Russ Bates, Sam Blackwell, Jason Yim, Olaf Ronneberger, Sebastian Bodenstein, Michal Zielinski, Alex Bridgland, Anna Potapenko, Andrew Cowie, Kathryn Tunyasuvunakool, Rishub Jain, Ellen Clancy, Pushmeet Kohli, John Jumper, and Demis Hassabis. Protein complex prediction with alphafold-multimer. bioRxiv, 2021.

[15] Diederik P. Kingma and Jimmy Ba. Adam: A method for stochastic optimization. arXiv preprint 1412.6980, 2014.

[16] Amelia Villegas-Morcillo, Louis Robinson, Arthur Flajolet, and Thomas D Barrett. ManyFold: an efficient and flexible library for training and validating protein folding models. Bioinformatics, 39(1):btac773, 12 2022.

[17] Valerio Mariani, Marco Biasini, Alessandro Barbato, and Torsten Schwede. lDDT: a local superposition-free score for comparing protein structures and models using distance difference tests. Bioinformatics, 29(21):2722–2728, 2013.

[18] Yang Zhang and Jeffrey Skolnick. Scoring function for automated assessment of protein structure template quality. Proteins: Structure, Function, and Bioinformatics, 57(4):702–710, 2004.

[19] Helen M. Berman, John Westbrook, Zukang Feng, Gary Gilliland, T. N. Bhat, Helge Weissig, Ilya N. Shindyalov, and Philip E. Bourne. The Protein Data Bank. Nucleic Acids Research, 28 (1):235–242, 2000.

[20] Xavier Robin, Juergen Haas, Rafal Gumienny, Anna Smolinski, Gerardo Tauriello, and Torsten Schwede. Continuous Automated Model EvaluatiOn (CAMEO)—Perspectives on the future of fully automated evaluation of structure prediction methods. Proteins: Structure, Function, and Bioinformatics, 89(12):1977–1986, 2021.

[21] Andriy Kryshtafovych, Torsten Schwede, Maya Topf, Krzysztof Fidelis, and John Moult. Critical assessment of methods of protein structure prediction (CASP)—Round XIV. Proteins: Structure, Function, and Bioinformatics, 89(12):1607–1617, 2021.

[22] Baris E. Suzek, Yuqi Wang, Hongzhan Huang, Peter B. McGarvey, Cathy H. Wu, and the UniProt Consortium. UniRef clusters: a comprehensive and scalable alternative for improving sequence similarity searches. Bioinformatics, 2014.

